# The Late Positive Event-Related Potential Component is Time-Locked to the Decision in Recognition Memory Tasks

**DOI:** 10.1101/2023.12.19.572476

**Authors:** Jie Sun, Adam F. Osth, Daniel Feuerriegel

## Abstract

Two event-related potential (ERP) components are commonly observed in recognition memory tasks: the Frontal Negativity (FN400) and the Late Positive Component (LPC). These components are widely interpreted as neural correlates of familiarity and recollection, respectively. However, the interpretation of LPC effects is complicated by inconsistent results regarding the timing of ERP amplitude differences. There are also mixed findings regarding how LPC amplitudes covary with decision confidence. Critically, LPC effects have almost always been measured using fixed time windows relative to memory probe stimulus onset, yet it has not been determined whether LPC effects are time-locked to the stimulus or the recognition memory decision. To investigate this, we analysed a large (n=l32) existing dataset recorded during recognition memory tasks with post-decisional confidence ratings. We used ERP deconvolution to disentangle contributions to LPC effects (defined as differences between hits and correct rejections) that were time-locked to either the stimulus or the vocal old/new response. We identified a left-lateralised parietal LPC effect that was time-locked to the vocal response rather than probe stimulus onset. We also isolated a response-locked, midline parietal ERP correlate of confidence that influenced measures of LPC amplitudes at left parietal electrodes. Our findings demonstrate that, contrary to widespread assumptions, the LPC effect is time-locked to the recognition memory decision and is best measured using response-locked ERPs. By extension, differences in response time distributions across conditions of interest may lead to substantial measurement biases when analysing stimulus-locked ERPs. Our findings highlight important confounding factors that further complicate the interpretation of existing stimulus-locked LPC effects as neural correlates of recollection. We recommend that future studies adopt our analytic approach to better isolate LPC effects and their sensitivity to manipulations in recognition memory tasks.

## 1. Introduction

Recognition memory refers to the ability to determine whether an item has been previously encountered. This allows us to discriminate between novel and familiar objects in our environment, which in turn underpins a range of cognitive functions such as learning and navigation based on familiar landmarks. Recognition memory has been extensively studied using mathematical models of retrieval and decision-making (Clark & Gronlund, 1996; Osth et al., 2018; Shiffrin & Steyvers, 1997) as well as neuroimaging techniques (Rugg & Curran, 2007; Rugg et al., 1999).

The number of distinct cognitive processes underlying recognition memory decisions has been a topic of ongoing debate. Advocates of dual process theories propose two distinct cognitive processes that support recognition memory decisions: recollection and familiarity (Atkinson & Juola, 1973; Diana et al., 2006; Wixted, 2007; Yonelinas, 1994). Recollection is proposed to support recognition and involves retrieving specific details about previously encountered items. It is conventionally considered an all-or-none, ‘high-threshold’ process that can discretely fail (but see Mickes et al., 2011; Onyper et al., 2010; Wixted, 2007). When recollection fails, observers must instead rely on processes that estimate a degree of familiarity. These processes are assumed to occur at earlier latencies than recollection. They support recognition based on a sense of knowing, but without retrieving source information or contextual details that were present during encoding. One’s degree of familiarity has been conceptualized as a continuous variable that covaries with recognition accuracy. Alternatively, computational models derived from single process theories, such as unequal-variance signal detection models (Ratcliff et al., 1992; Slotnick & Dodson, 2005) and global matching models (e.g., Cox & Shiffrin, 2017; Dennis & Humphreys, 2001; Gillund & Shiffrin, 1984; Osth & Dennis, 2015; Shiffrin & Steyvers, 1997), specify that recognition memory decisions are based on internally-derived estimates of item familiarity or another strength-of-memory variable.

### 1.1. Neural Correlates of Familiarity and Recollection

Using electroencephalography (EEG), researchers have investigated the neural correlates of recognition memory performance using single and dual process frameworks. Two event-related potential (ERP) components have been found to be related to recognition performance, namely the Frontal Negativity (termed the FN400) and Late Positive Component (LPC; Curran, 2000; Rugg & Curran, 2007; Rugg & Yonelinas, 2003). The FN400 is defined as a negative-going ERP component at frontal electrodes that peaks around 400 ms after the onset of memory probe stimulus. More positive-going FN400 amplitudes have been observed in trials with hits (i.e., successful recognition of previously seen items) compared to correct rejections following novel items (e.g., Curran & Hancock, 2007). The LPC is commonly measured at left parietal electrodes with a positive peak occurring between 400-800 ms after memory probe onset. More positive-going LPC amplitudes are observed in trials with hits as compared to correct rejections (e.g., Addante et al., 2012). Within the dual process framework, FN400 effects have been interpreted as reflecting differences in familiarity, whereas LPC effects have be interpreted as reflecting differences in the proportion of trials involving successful recollection (e.g., Curran & Cleary, 2003; Friedman & Johnson Jr, 2000; Rugg & Curran, 2007).

This dual process interpretation is based on findings suggesting a double dissociation between FN400 and LPC effects. First, the amplitude of the FN400, but not the LPC, was found to vary with manipulations of familiarity, operationalised as the stimulus similarity across targets and lures. FN400 amplitudes have been reported as more positive-going for false alarms to lures that were similar to targets, as compared to lures that were dissimilar, in experiments presenting words (Curran, 2000), pictures (Curran & Cleary, 2003) and faces (Nessler et al., 2005). By contrast, differences in FN400 amplitudes were not observed across conditions that are thought to induce different rates of successful recollection (e.g., Chan & McDermott, 2007; Rugg et al., 1998). On the other hand, LPC amplitudes have been reported as more positive-going in trials with ‘remember’ as compared to ‘know’ responses (Sarah & Rugg, 2010; Woodruff et al., 2006), which are argued to reflect the decisions based on recollection and familiarity, respectively (Tulving, 1985; Yonelinas, 1994). Further, more positive LPC amplitudes were found for words processed semantically and elaborated in a sentence during encoding (Rugg et al., 1998), repeated exposure to pseudowords as opposed to a single presentation during encoding (Bermúdez-Margaretto et al., 2015), and for stimuli encoded under focused as compared to divided attention (Curran, 2004). These conditions are thought to increase the likelihood of recollection but minimally affect familiarity (Gardiner & Parkin, 1990; Rugg et al., 1998). In addition, FN400 amplitudes were found to continuously vary with one’s degree of decision confidence (Woodruff et al., 2006; Sarah & Rugg, 2010) while increases in LPC amplitudes were only observed for high-confidence old responses (Addante et al., 2012). Collectively, these findings support the notion that the FN400 and LPC components index two distinct cognitive processes, whereby the FN400 is consistent with a continuous familiarity process while the LPC aligns with a high-threshold recollection process.

However, others have claimed that the remember-know paradigm does not selectively influence familiarity and recollection (Donaldson, 1996; Dunn, 2004; Rotello et al., 2005), in which each type of response does not indicate a distinct process but rather reflect one common process on a continuum. Further, studies have investigated the dimensionality of recognition memory using state-trace analyses, a method to determine the latent structures of the effects across tasks/conditions that is superior to traditional dissociation methods (Newell & Dunn, 2008). Using this method, little evidence has been found for two distinct processes in the remember-know paradigm (Dunn, 2008). By using state trace analysis to investigate changes in FN400 and LPC amplitude under different degrees of attention, Freeman et al. (2010) also concluded that the results were consistent with a single process rather than a dual process interpretation of recognition memory.

### 1.2. LPC Amplitude Covariations with Decision Confidence

In addition, inconsistent findings have been reported regarding the relationships between these ERP components and decision confidence. Using a remember-know design, high confidence “know” responses - which are theorised to rely on familiarity - showed greater LPC amplitude than low confidence “remember” responses that are presumed to index recollection (Brezis et al., 2017). Further, LPC amplitudes were found to vary with different degrees of decision confidence for both hit and correct rejection responses (Curran & Hancock, 2007; Rubin et al., 1999; Woroch & Gonsalves, 2010), instead of being exclusively associated with high confidence hits as would be expected if the LPC indexes recollection. These findings are inconsistent with a high-threshold recollection process assumed in classic dual process models (Yonelinas, 1994), but are consistent with notions of recollection being a continuous process, as well as single process theories of recognition memory (Ratcliff et al., 1992, Slotnick & Dodson, 2005).

Studies investigating this inconsistency having identified a separate confidence signal in parietal regions that overlaps with the LPC (Curran & Hancock, 2007; Woodruff et al., 2006) and may contribute to graded changes in measured LPC amplitudes across confidence ratings. However, as recollection is assumed to cause high decision confidence (Yonelinas, 1994), it becomes difficult to determine whether LPC effects are related to recollection or merely high confidence responses due to large-amplitude, confidence-related signals. Therefore, understanding how confidence-related signals influence LPC measures is crucial for concluding whether the LPC effect indexes a high-threshold recollection process. So far, findings seem to diverge on the temporal and spatial characteristics of this confidence-related component (time window of 500 - 800 ms, bilateral parietal electrodes, Woodruff et al., 2006; 700 - 800 ms, central electrodes, Curran & Hancock, 2007).

### 1.3. Stimulus- and Response-Locked Contributions to LPC Effects

In addition to the conceptual issues described above, there is also the issue of how FN400 and LPC effects are most appropriately measured. Notably, LPC mean amplitude measurement windows have varied markedly across studies (Addante et al., 2012; Brezis et al., 2017; Mograss et al., 2009; Woroch & Gonsalves, 2010; Zhang et al., 2015). In some studies, the latency of the LPC was suggested to be earlier for remember than know responses (Spencer et al., 2000), and earlier for primed probe words than unprimed words (Li et al., 2017; Woollams et al., 2008). The issue of ERP component latency changes across conditions is not uncommon in decision-making research (Ouyang et al., 2017), and leads to confounds when contrasting conditions with different component latencies. For example, if an ERP component occurs at different times for separate conditions and/or participant populations (e.g., due to age differences, Jaworska et al., 2020), there will likely be mean amplitude differences within a defined time window between those conditions/groups, even if the ERP components are identical in amplitude. However, the reasons behind the inconsistent latency of LPC effects has not been systematically investigated, and studies have frequently adopted *a priori* time window assumptions for EEG studies of recognition memory (Hubbard et al., 2019; Wynn et al., 2020; Yang et al., 2019).

A simple explanation for this latency variability is that the LPC is time-locked to the recognition memory decision and associated motor response. For example, another ERP component—the central parietal positivity (CPP), a correlate of evidence accumulation and confidence in perceptual decision-making tasks (Grogan et al., 2023; Kelly & O’Connell, 2013; Ko et al., 2023; O’Connell et al., 2012) — peaks just prior to the time of the response. Consequently, the CPP peaks at longer latencies from stimulus onset in trials with slower response times (RTs, Feuerriegel et al., 2022; Kelly & O’Connell, 2013; O’Connell et al., 2012). Similarly, if the LPC is time-locked to the response, the latency differences observed in stimulus-locked data could be explained by different RT distributions across conditions of interest. If assuming that LPC effects index recollection, then successful recollection could rapidly lead to a memory decision immediately followed by a motor response, making the LPC effect time-locked to the response rather than probe stimulus onset.

Indeed, there is tentative evidence that LPC latency increases with longer reaction times (Li et al., 2017; Marsh, 1975). Further, a response-locked, left-lateralised parietal old/new effect in recognition memory tasks has been observed using response-locked data (Kayser et al., 2007; Kayser et al., 2009). While initial evidence seems to indicate that LPC could be time-locked to the response, this is yet to be clearly determined. First, latency changes in stimulus-locked data do not provide direct evidence that LPC is time-locked to the response. Second, response-locked ERPs in trials with faster RTs occur closer to stimulus onset and therefore overlap more with ERP waveforms that are time-locked to the stimulus. This can create biases in EEG amplitude measures when contrasting conditions with different RT distributions.

### 1.4. Disentangling Different Contributions to FN400 and LPC Effects

To determine the timing of LPC effects and overcome potential confounds from neural signals that are time-locked to the stimulus onset, we used a method that decomposes EEG signals into subcomponents called Residual Iteration Decomposition (RIDE, Ouyang et al., 2011; Ouyang et al., 2015). This method was introduced to separate subcomponents of ERPs that are time locked to different events in each trial, such as the onset of a stimulus and the behavioural response (respectively termed stimulus-locked and response-locked subcomponents). This is achieved by an iterative estimation process that estimates ERP waveforms that are closely time-locked to the stimulus across trials, as well as waveforms that are variable in time or time-locked to the response. By comparing the stimulus-locked subcomponent across hit and correct rejection conditions, we could assess ERP effects that are closely time-locked to probe stimulus onset. By subtracting the stimulus-locked subcomponents from the single-trial ERP data for each condition, we could separately estimate the magnitudes of ERP effects that are variable in time, such as those that are time-locked to the response (as done by Steinemann et al., 2018). This approach has been successfully used to dissociate stimulus- and response-locked ERP effects that are typically intermixed in speeded decision-making tasks (e.g., Ouyang et al., 2017; Verleger et al., 2015; Verleger et al., 2014).

In order to resolve debates around the functional significance of FN400 and LPC effects, it is critical to first determine how these components should be most appropriately measured. In this study, we investigated whether the LPC effect is time-locked to the memory probe stimulus or the recognition memory decision. We analysed a large (n=!32) existing EEG dataset collected by Weidemann and Kahana (2019) that was recorded during a recognition memory task involving lists of words. This dataset also included confidence ratings for each trial, which allowed us to additionally characterise the neural correlates of decision confidence in this recognition memory task. To foreshadow our results, we found that the LPC effect was closely time-locked to the response rather than memory probe onset, contrary to widely held assumptions. We also identified an additional, response-locked parietal correlate of confidence that had a distinct topography from LPC effects occurring over the same time window. Our results have direct implications for the interpretation of existing LPC effects measured using stimulus-locked ERPs, as well as the broader debate around the interpretation of LPC effects based on this evidence.

## 2. Method

### 2.1. Participants

We analysed an online archival dataset that is described in Weidemann and Kahana (2019). The dataset contains task performance and EEG data from 132 participants (age range 17-30, mean = 22.1, SD = .3), which is a subset of data from the Penn Electrophysiology of Encoding and Retrieval Study (PEERS) including experiments 1, 2 and 3 (Kahana et al., 2023). This dataset included test phase data only. Participants consisted of students and staff at the University of Pennsylvania, Drexel University, Rowan University, Temple University, University of the Arts, and the University of the Sciences. For each participant, 20 sessions of EEG data were collected (average of 3,645 trials per participant included in analyses). A detailed description of participant characteristics can be found in Weidemann and Kahana (2016). The collection of this dataset was approved by the Institutional Review Board of the University of Pennsylvania.

### 2.2. Paradigm

Task performance and EEG data were recorded during recognition memory experiments. For each experiment, different variants of encoding and recall tasks were used (for a detailed description, see Kahana et al., 2023). Therefore, some variation will present in the tasks related to the recognition performance across sessions and subsets of the participants.

The structure of the experiment is shown in Figure 1. In each session participants were instructed to study either 12 or 16 lists of 16 words each. Each word was presented for 3 s with an interstimulus interval ranging between 0.8 and 1.2 s. During this study phase, participants were additionally required to make either a size judgement (“Will this item fit into a shoebox?”) or an animacy judgement (“Does this word refer to something living or not living?”) about the presented word. At the end of each list, participants were instructed to freely recall the items in the study list for 75 s (i.e., an immediate recall task). For a randomly determined half of the sessions per participant, an additional final free recall task was included in which participants were given 5 minutes to freely recall any items studied in this session.

**Figure 1.**
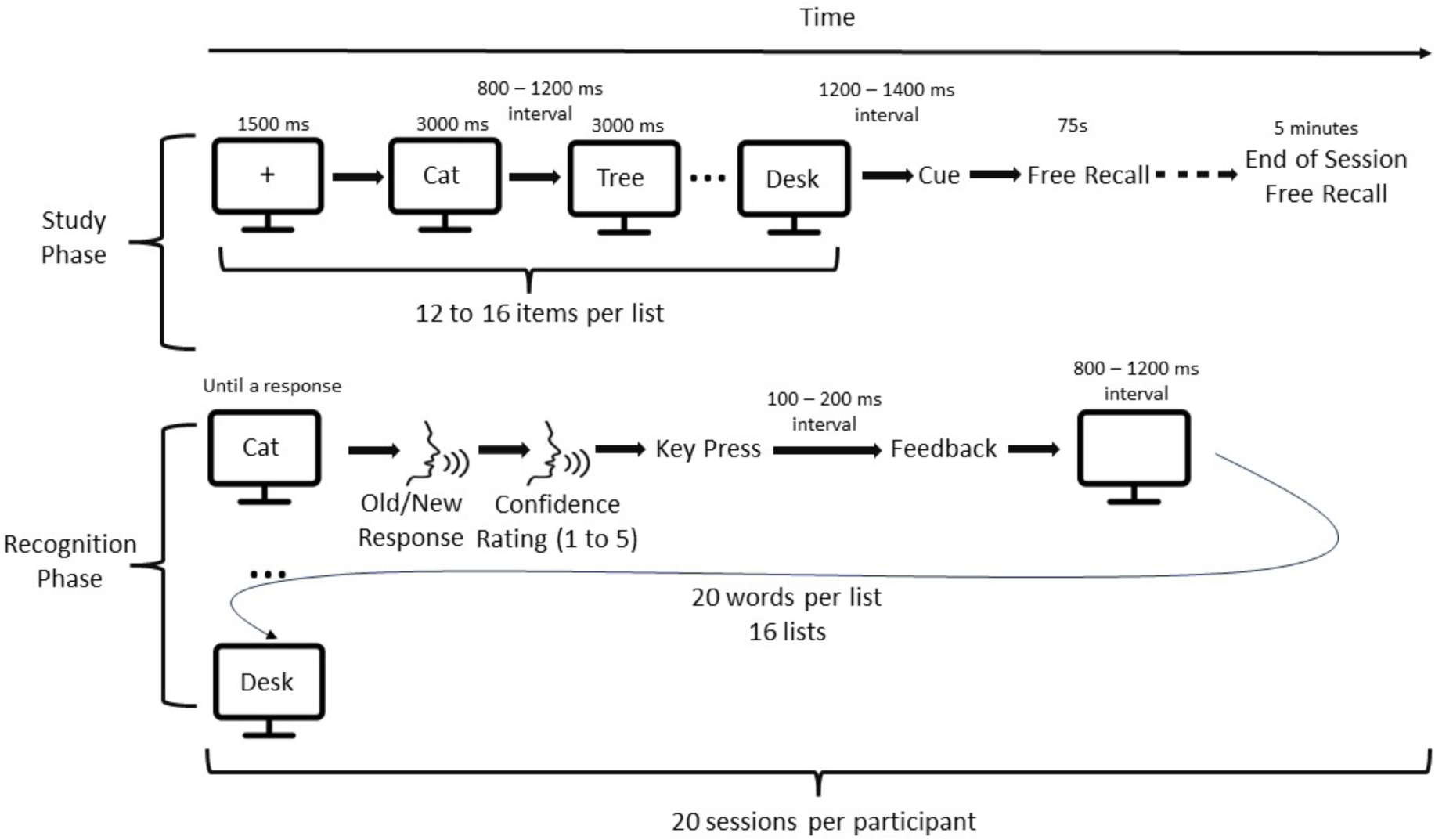
Experiment structure for the dataset described in Weidemann and Kahana (2019).

After viewing the study lists and completing the immediate recall task, participants were instructed to complete a recognition memory task. The recognition task contained 320 probe items of which either 80%, 75%, 62.5% or 50% were targets (varying across sessions) and the rest were lures. Participants were required to respond verbally (“Pess” for “yes” and “Po” for “no”) to indicate whether they had seen the item in the study lists. This was followed by an immediate verbal confidence judgement ranging from 1 to 5, indicating low to high decision confidence regarding the old/new decision in that trial. Please note that this differs from experiments whereby participants provide their choice and confidence simultaneously (e.g., ’confident old’, Curran & Hancock, 2007; Summerfield & Mangels, 2005; Woroch & Gonsalves, 2010). Feedback on the recognition judgement was provided after an interval of 100-200 ms following the confidence report. The next probe item appeared after an 800-1200 ms interval (varied across trials).

### 2.3. Stimuli

The complete set of stimuli included 1,638 words selected for all the PEERS experiments (available at https://memory.psych.upenn.edu/Word_Pools). Word lengths ranged from 2 to 12 letters (median = 6), and word frequencies (per million) ranged from .06 to 5,247 (median = 7.5), according to SUBTLEXus database (Brysbaert & New, 2009). Given the animacy and size judgements implemented in the study phase, all words were selected based on whether they had a clear meaning with respect to animacy and size.

### 2.4. EEG Data Acquisition

High density EEG data were recorded using the 129 channel EGI system with a sampling rate of 500 Hz. Channel Cz was used a reference during the recording, and all recordings were re-referenced to an average reference offline. A high-pass filter at 0.1 Hz was applied during recording. The data described here was from the recognition memory task phase of each session. Study phase data was not available from the online repository.

### 2.5. EEG Data Processing

For the archival dataset, twenty-six electrodes placed on the face were excluded from data processing and analysis, leaving 103 channels in total. EEG data were segmented from -2,200 ms prior to the target/lure onset until 2,000 ms after the response. Trials with response times faster than 300 ms and slower than 3,000 ms were excluded (as done in Weidemann & Kahana, 2019).

After receiving this dataset with the processing steps applied above, we further processed the data using EEGLAB (v 2021.1 Delorme & Makeig, 2004) running in MATLAB (The Mathworks). Code used for data processing and analysis is available at https://osf.io/kur25. Data processing was done for each session separately. We shortened the epochs to range from -1,000 ms to 2,200 ms relative to target/lure onset and excluded trials with RTs longer than 2 seconds. Data were then low-pass filtered at 30 Hz (EEGLAB Basic Finite Impulse Response Filter New, zero-phase, -6 dB cutoff frequency 33.75 Hz, transition band width 7.5 Hz) and re-referenced using a linked mastoids reference. Channels were marked as excessively noisy if they had amplitudes of ± 500 µV for at least 20% of all the trials in one session, and if the channel values greatly deviated from the centre of the distribution of all channel values (> 5 standard deviations). These channels were excluded from further processing steps and were later interpolated.

Trials with EEG amplitudes exceeding ± 500 µV were rejected. Following this, an Independent Component Analysis (ICA) was done for each session (RunICA extended algorithm, Jung et al., 2000). We then identified and subtracted components associated with ocular artefacts using ICLabel (Pion-Tonachini et al., 2019). Components identified as eye artefacts with over 85% probability were subtracted. We then interpolated the excessively noisy channels using spherical spline interpolation. Segments were then baseline-corrected using the 200 ms window preceding stimulus onset. Trials with amplitudes exceeding ± 200 µV were removed.

We then aggregated EEG data from all sessions within each participant. The stimulus-locked segments used for analyses were from -1,000 to 2,200 ms relative to stimulus onset, Response-locked segments were created from -1,300 to 200 ms relative to the onset of the vocal response. Processed data used for analyses are available at https://osf.io/kur25.

### 2.6. Deconvolving Stimulus- and Response-Locked Contributions to ERPs

We used the RIDE toolbox (Ouyang et al., 2011; Ouyang et al., 2015) to separate subcomponents of the ERP signals that are time-locked to either stimulus onset (termed the stimulus-locked subcomponent) or the response (termed the response-locked subcomponent). RIDE achieves this by using an iterative algorithm that produces estimates of each deconvoluted subcomponent within each iteration until the estimates converge. The estimates were produced within pre-defined windows for each subcomponent. To cover the full range of pre-defined time windows for the FN400 and LPC components, we used a 0 - 800 ms time window relative to stimulus onset for the stimulus-locked subcomponent and -600 to 200 ms relative to the onset of the vocal response for the response-locked subcomponent. The input epochs ranged from - 200 ms to 2200 ms relative to the stimulus onset. The waveforms of each subcomponent were separately derived with the epoch length equal to the input epoch. For the response-locked data, we additionally subtracted the estimated stimulus-locked subcomponent for each participant before realigning stimulus-locked data to the response in each trial (as done by Steinemann et al., 2018). The above procedures were done for each participant separately using data pooled across testing sessions.

### 2.7. Characterising FN400 and LPC Effects in Stimulus- and Response-Locked Data

We characterised the time-courses of FN400 effects (at electrode Fz) and LPC effects (at electrode P3) by comparing ERP amplitudes across trials with hits and correct rejections. We performed mass univariate analyses with cluster-based permutation tests (Groppe et al., 2011; Maris & Oostenveld, 2007) as implemented in the Decision Decoding Toolbox (Bode et al., 2019) to correct for multiple comparisons. We first compared ERP amplitudes at each timepoint from stimulus or response onset using paired-samples t tests. Neighbouring timepoints with statistically significant differences (p < .01) were grouped into clusters. The mass of each cluster was calculated as the sum of t values within that cluster. This procedure was then repeated for 10,000 permutation samples whereby condition labels were switched for a subset of participants. The maximum cluster mass for each permutation sample formed the empirical null distribution. Clusters in the original dataset with larger masses than the 97.5^th^ percentile of the null distribution (corresponding to an alpha level of 0.05, two tailed) were marked as statistically significant.

This was first done for probe stimulus-locked ERPs (spanning -200 to 800 ms from stimulus onset) to determine whether we could replicate typical FN400 and LPC effects (e.g., Rugg & Curran, 2007). For ERPs at electrode P3 (corresponding to the LPC effect) we repeated these analyses using the stimulus-locked subcomponent isolated using the RIDE method. We also performed these analyses using stimulus-locked ERPs whereby the stimulus-locked subcomponent was subtracted from each trial. This allowed us to quantify the contributions of ERP effects that were not strictly time-locked to probe stimulus onset (e.g., time-varying or response-locked effects) on stimulus-locked ERPs that have been typically analysed in prior work. We also compared ERPs across hits and correct rejections using response-locked data at electrode P3 with stimulus-locked subcomponent subtracted (epochs spanning -600 to 200 ms relative to the onset of the vocal response).

### 2.8. Current Source Density (CSD) Transformation

Given the reported temporal overlap between FN400 and LPC effects, it is likely that effects at parietal electrodes (e.g., P3) may influence ERP measurements at frontal channels (e.g., Fz) due to the broad spread of electrical brain potentials across the scalp (known as volume conduction). As both ERP components are defined by contrasting hits and correct rejections, we were concerned about effects of signal spread from parietal to frontal scalp electrodes. To address this, we derived Current Source Density (CSD) transformed ERPs to test for FN400 effects using mass-univariate analyses while minimising volume conduction from parietal sources (Nicholson & Freeman, 1975). This was done using the CSD toolbox in MATLAB (m-constant = 4, λ = 0.00001, Kayser & Tenke, 2006). The method takes spatial derivatives of the scalp EEG data and reduces the point spread from each channel.

### 2.9. Assessing ERP Correlates of Confidence

We additionally characterised ERP correlates of decision confidence during the time window of the response-locked LPC effects identified using mass-univariate analyses. For each electrode, and for trials with hits and correct rejections separately, we fit linear mixed effects models using the confidence ratings in each trial to predict single-trial mean ERP amplitudes across the time window of-200 to 0 ms relative to the old/new vocal response. The time window was selected to avoid contamination from the speech artefacts associated with the verbal old/new responses (as noted in Kahana et al., 2023). We also included random intercepts and slopes across participants. The model equation is as follows:

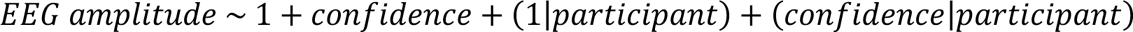

The obtained regression coefficients (betas) for the predictor of confidence at each electrode indicated the strength and direction of the association between the EEG amplitude and the degree of confidence. To test for effects of confidence on response-locked LPC amplitudes, we derived p-values corresponding to the fixed effect of confidence at electrode P3 for hits and correct rejection trials separately.

## 3. Results

### 3.1. Task Performance

Participants showed good recognition memory performance. For the set of trials included for EEG analyses, participants achieved a 91.1% hit rate on average (range = 76.5 - 98.7%, SD = 4.5%) and a false alarm rate of 14.2% (range = 1.7 - 45.2%, SD = 10.4%). Mean RTs for hits (mean = 756 ms, range = 479 - 1161 ms, SD = 104 ms) were faster than those for correct rejections (mean = 935 ms, range = 590 - 1382 ms, SD = 122 ms), ř(l3l) = -36.9, *p* < .001, consistent with previous studies (Maratos et al., 2000; Ratcliff & Murdock, 1976; Wilding & Rugg, 1996).

### 3.2. Characterising FN400 Effects

At electrode Fz we identified a time window spanning approximately 400-640 ms from stimulus onset whereby ERP amplitudes were more positive-going for hits as compared to correct rejections (Figure 2A). This resembles the typical FN400 effect, although the time window is later than reported in some previous work (Curran & Hancock, 2007; Leynes et al., 2017; Rugg & Curran, 2007).

**Figure 2.**
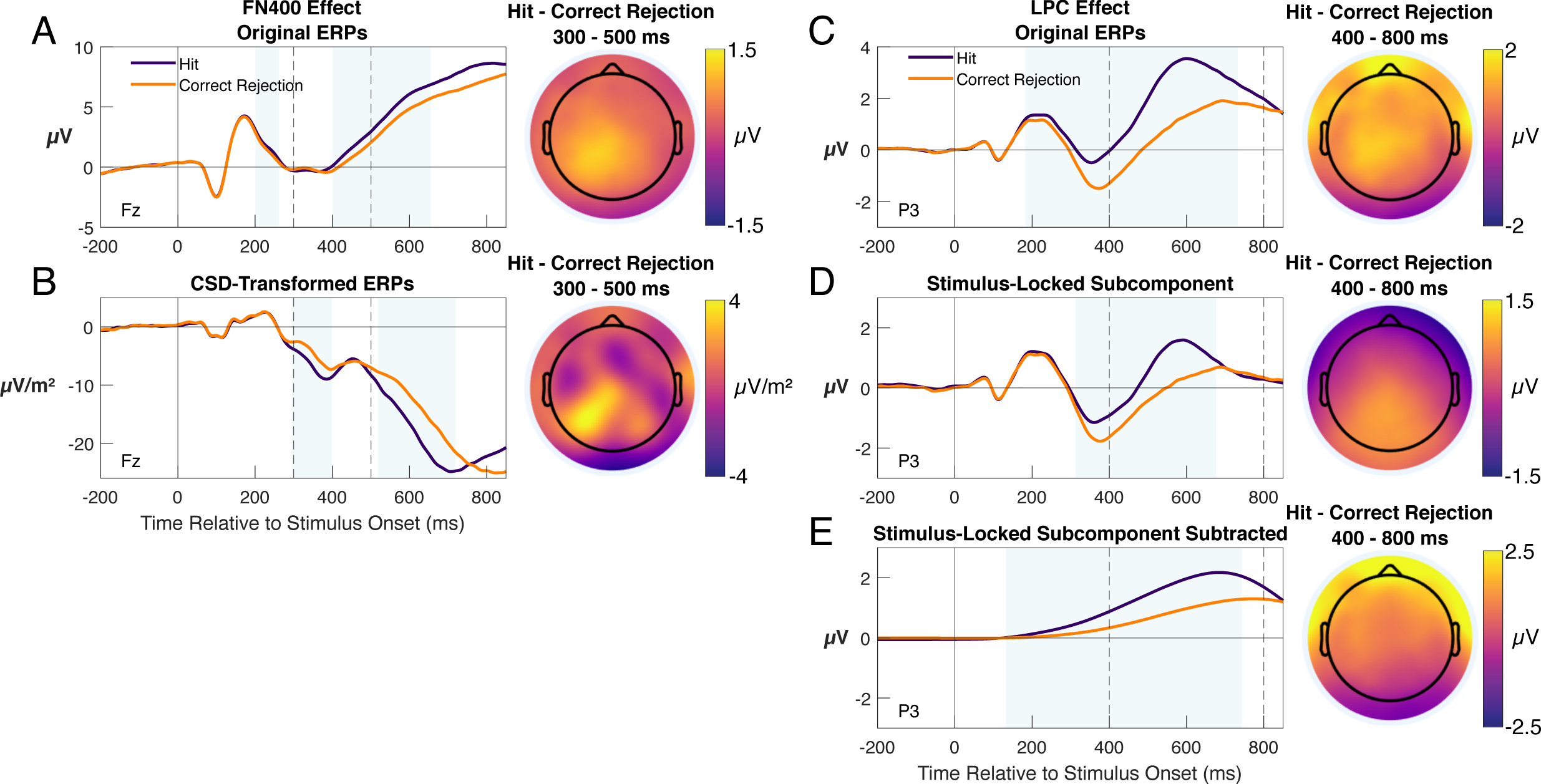
ERPs for hits and correct rejections at electrodes Fz and P3, corresponding to the FN400 and LPC components. A) ERPs for hits and correct rejections in the original data at electrode Fz and the scalp map of amplitude differences averaged over 300 - 500 ms from target/lure onset. B) CSD-transformed ERPs at Fz. C) ERPs for hit and correct rejection trials in the original data at electrode P3 and the scalp map of amplitude differences averaged over 400 - 800 ms from target/lure onset. D) ERPs for the stimulus-locked subcomponent at electrode P3. E) ERPs for the stimulus-locked data at electrode P3 after the subtracting stimulus-locked subcomponent. Dashed lines indicate the conventional time windows used to measure the FN400 and LPC components in previous studies. Blue-shaded areas denote statistically significant differences identified using mass univariate analyses.

Notably, the scalp distribution of this effect (Figure 2A) did not show a clear frontal locus. Instead, amplitude differences were largest over left parietal channels that are typically used to measure the LPC over similar time windows. This suggests that more positive-going ERP amplitudes at frontal channels may have arisen from posterior parietal effects spreading across the scalp to Fz, rather than a genuine frontal effect (i.e., volume conduction, for similar effects spanning frontal and parietal channels see Feuerriegel et al., 2022). To better isolate frontal and parietal sources, we performed the same analyses using CSD-transformed data. We did not observe the typical FN400 effect at Fz; statistically significant ERP differences were in the opposite direction (i.e., more positive-going for correct rejections, Figure 2B). By contrast, the LPC effect with a left parietal locus was clearly visible. As we did not identify an FN400 effect with a clear frontal locus, we did not perform further analyses using deconvolved ERPs at electrode Fz.

### 3.3. Characterising LPC Effects

At electrode P3 we observed more positive-going ERPs for hits compared to correct rejections (i.e., a typical LPC effect, Figure 2C) consistent with previous work (Curran, 2000; Rugg & Yonelinas, 2003, Rugg & Curran, 2007). This left lateralized effect was observed over a wide time range (approximately 200-730 ms) which overlapped with standard definitions of LPC effects spanning 400-800 ms after target/lure onset (Curran & Cleary, 2003; Friedman & Johnson Jr, 2000; Rugg & Curran, 2007).

When using RIDE to deconvolve stimulus-locked and non-stimulus-locked EEG signals, we observed a similar effect for the extracted stimulus-locked subcomponent at electrode P3 (spanning 310 to 690 ms, Figure 2D). However, the scalp topography of this stimulus-locked effect was most prominent at midline parietal areas, differing from the left-parietal distribution of the typical LPC. By contrast, comparisons between hits and correct rejections using data with the stimulus-locked subcomponent subtracted (Figure 2E) showed a clear left-lateralised parietal effect between 130 and 740 ms. Despite both stimulus-locked and non stimulus-locked effects contributing substantially to the original LPC effect, only the non stimulus-locked subcomponent showed a topography that was left-lateralised (in line with the definition of LPC effect, Friedman & Johnson Jr, 2000; Rugg & Curran, 2007).

To verify whether the left-lateralised LPC effects were indeed time-locked to the response, we subtracted the stimulus-locked subcomponent of the ERP (derived separately for each participant and each hit/correct rejection condition) from each stimulus-locked trial of EEG data (as done by Steinemann et al., 2018). We then derived epochs that were time-locked to the onset of the vocal response. This allowed us to better isolate any response-locked LPC effects that were independent of the co-occurring stimulus-locked parietal effect identified above.

We observed more positive-going amplitudes for hits compared to correct rejections immediately preceding the vocal response (-180 to -50 ms, Figure 3A) and from 40 ms after the response. The scalp map in Figure 3B displays a left-lateralised topography as is typical of the LPC effect found in previous work.

**Figure 3.**
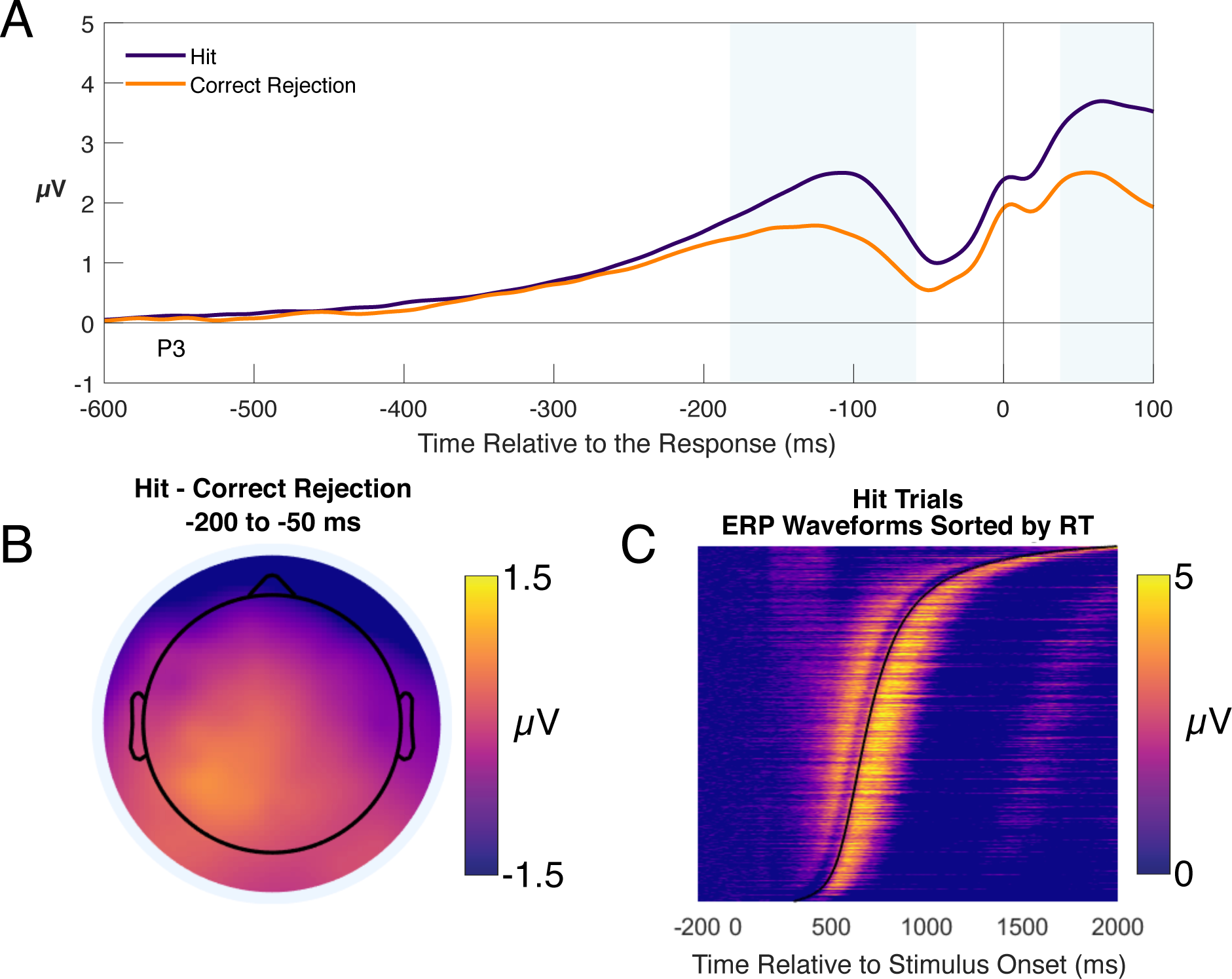
EEG data for hits and correct rejections following subtraction of the stimulus-locked ERP subcomponent. A) Response-locked ERPs at electrode P3. Blue-shaded areas mark statistically significant differences identified using mass univariate analyses. B) Scalp map of hit and correct rejection differences over the -200 to -50 ms pre-response time window. C) Stimulus-locked ERPs with stimulus-locked subcomponents subtracted (pooled across participants) for hit responses at electrode P3. Trials are arranged by response time (indicated by the curved black line). Data were smoothed using a sliding Gaussian window spanning 500 trials.

The response-locked nature of this effect can also be observed by plotting single-trial ERP amplitudes for hit responses, pooled across participants and sorted from fastest to slowest RT (Figure 3C, as done by O’Connell et al., 2012; Feuerriegel et al., 2022). This plot displays prominent, positive-going amplitudes immediately preceding the time of the response, followed by another positive-going response immediately after the response is initiated.

Taken together, these results show that there are two dissociable contributions to the LPC effects in our original stimulus-locked ERPs: one with a midline parietal topography that is time-locked to stimulus onset (Figure 2D) and another with a left-lateralised topography that is time-locked to the response (Figure 3). Notably, the stimulus-locked ERPs are also influenced by effects that occur after the recognition memory decision has been made in each trial (corresponding to post-response ERP differences).

### 3.4. Response-Locked ERP Correlates of Decision Confidence

Having identified a response-locked LPC effect, we additionally assessed whether response-locked ERPs at electrode P3 also covary with decision confidence. As parietal ERP correlates of confidence have been identified in perceptual decision-making tasks (e.g., Feuerriegel et al., 2022; Grogan et al., 2023; Ko et al., 2023) we also sought to determine whether the scalp distributions of confidence effects were distinct from the left-lateralised topography of the LPC effect. For these analyses we used response-locked single-trial ERPs with the stimulus-locked subcomponent subtracted. Please note that the dataset included confidence ratings ranging from 1 to 5 (from lowest to highest) for either ‘old’ or ‘new’ responses. For comparability with existing work that investigated effects of confidence on LPC amplitudes (e.g., Addante et al., 2012; Woodruff et al., 2006; Woroch & Gonsalves, 2010), we have also plotted ERPs for hits followed by high (5) and lower (T4) confidence ratings, as well as ERPs for correct rejections, in the Supplementary Material.

Results from mixed effect models showed that pre-response amplitudes at electrode P3 significantly covaried with confidence for both hits *(β* = .30, 95% CI = [.23, .36], *p* < .001, Figure 4A) and correct rejections *(β =* .11, 95% CI = [.02, .21], *p* = .019, Figure 4C). More positive-going amplitudes were observed in trials with higher confidence ratings during the -200 to 0 ms pre-response time window.

**Figure 4.**
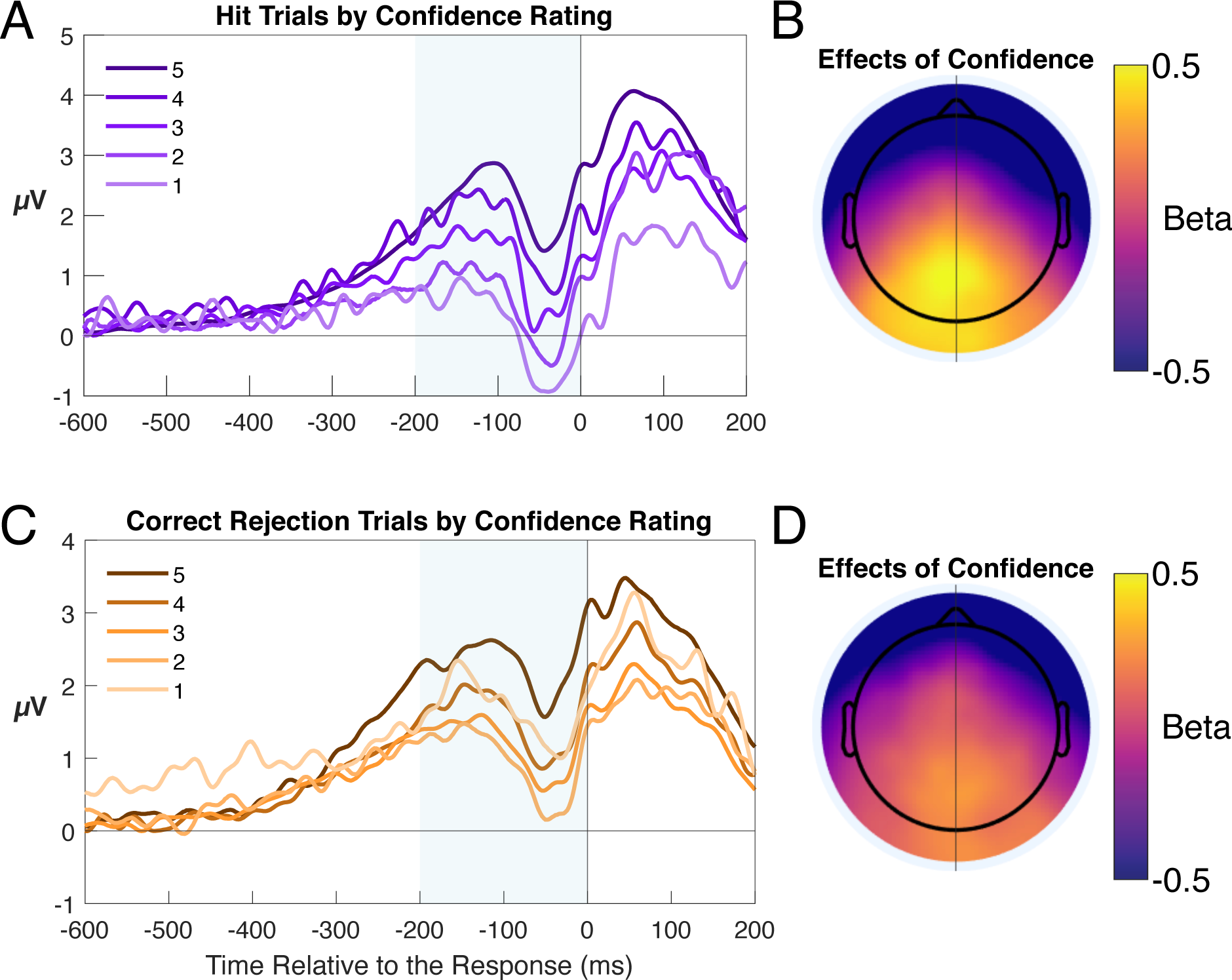
Response-locked ERPs by confidence rating after subtracting the stimulus-locked subcomponent from each trial. A) ERPs for hits at electrode P3. B) Scalp map of group-averaged regression slope coefficients (betas) from models using confidence to predict pre-response amplitude (averaged from -200 to 0 ms to response) for hit trials. C) ERPs for correct rejections. D) Scalp map of regression coefficients for correct rejection trials. Shading denotes the time windows used to calculate mean ERP amplitudes that were entered into the mixed effects models.

We additionally fit mixed effects models to pre-response ERP amplitudes at each electrode to derive a topography of associations with confidence (displayed in Figures 4B, D). Group-averaged beta values (indicating the slope of the association between confidence and amplitude) were focused over midline parietal channels, consistent with previous observations of a parietal confidence-related signal (Woodruff et al., 2006; Curran & Hancock, 2007). However, in contrast to those studies using stimulus-locked ERPs, our results indicate that this midline parietal confidence signal is time-locked to the response. Notably, this confidence-related effect temporally overlapped with the response-locked, left-lateralised LPC effect that was observed when contrasting hits and correct rejections (compare Figures 3B and 4B).

To summarise, we identified a response-locked, midline parietal ERP correlate of confidence in both hit and correct rejection trials, which was topographically distinct from the left-lateralised LPC effect but occurred over similar time windows relative to the response. This effect of confidence influenced ERP amplitudes at electrode P3, which is typically used to measure LPC effects.

## 4. Discussion

We systematically investigated the different signals that contribute to ERP correlates of recognition memory. By applying ERP deconvolution techniques to a large EEG dataset recorded during a typical recognition memory task, we aimed to isolate and quantify any stimulus- and response-locked signals that contribute to FN400 and LPC effects. We demonstrate that the canonical LPC effect (measured at left parietal electrodes) is time-locked to the recognition memory decision and associated motor response. This is contrary to the standard definition of this ERP component and widespread assumptions of a fixed component latency relative to memory probe onset. We additionally observed two other signals that influenced measures of LPC effects at left parietal channels: response-locked effects that extended into the post-response period (i.e., after the recognition memory decision had been made) and a stimulus-locked, midline parietal subcomponent. We also identified a response-locked neural correlate of decision confidence focused over midline parietal electrodes, which influences LPC amplitude measures at left parietal channels. Contrary to our expectations, we did not identify an FN400 effect with a clear source over frontal electrodes. Our findings show that multiple ERP signal sources can contribute to LPC effects measured using stimulus-locked ERPs, and that these are best disentangled using response-locked ERPs in combination with deconvolution methods. Our findings highlight important issues that complicate the interpretation of prior, stimulus-locked LPC effects in relation to recollection and decision confidence.

### 4.1. Response-Locked Contributions to LPC Effects

In our response-locked ERPs, we identified more positive-going amplitudes for hits compared to correct rejections immediately prior to the old/new vocal response in each trial. We observed a left parietal topography that is typical of the LPC effect. By contrast, the stimulus-locked ERP effects in our data were instead focused over midline parietal channels. This indicates that, although there are distinct sources that contribute to measures of LPC effects in stimulus-locked ERPs, the canonical, left-lateralised parietal LPC effect is time-locked to the recognition memory decision and associated motor response.

As RTs vary widely across trials in recognition memory tasks, this also means that the latency of LPC effects will also substantially vary across trials. This contrasts with widespread assumption that LPC effects occur within a fixed time window relative to stimulus onset (e.g., 400 - 800 ms, Rugg & Curran, 2007). However, our findings are congruent with existing observations of LPC effects with a varying latencies in stimulus-locked data (Li et al., 2017; Marsh, 1975) and LPC-like effects observed in response-locked ERPs (Kayser et al., 2007; Kayser et al., 2009).

This redefinition of LPC effects is practically and theoretically important. We propose that LPC amplitude measures using stimulus-locked ERPs are susceptible to sources of variance (and potentially bias) depending on the RT distributions for hits and correct rejections. For example, if RTs are faster for hits compared to correct rejections, as observed in behavioural (Ratcliff & Murdock, 1976; Wilding et al., 1996) and EEG studies (Glaser et al., 2012; Medrano et al., 2017; Rubin et al., 1999; Yu et al., 2022) as well as our own data, the LPC will peak earlier relative to stimulus onset for hits. This would produce apparent stimulus-locked amplitude differences that are partly due to shifts in ERP component latency (rather than amplitude). In addition, any changes in the distributions of RTs across conditions would also be expected to lead to different proportions of trials in which the LPC peak is captured within the canonical stimulus-locked mean amplitude time window (discussed in Feuerriegel et al., 2022). Although RT differences do not always occur across hits and correct rejections, in cases where they are observed (e.g., due to a bias toward ‘old’ responses) the interpretation of stimulus-locked LPC effects is complicated by potential changes in both component latency and amplitude.

Due to the complexity of these RT-related effects on ERP waveforms, we have refrained from making specific claims about existing studies here. However, we note that manipulations of familiarity and recollection in recognition memory tasks (e.g., depth of processing manipulations) are likely to also influence recognition RTs and lead to the issues described above. Re-analysis of existing datasets using ERP deconvolution techniques could determine how, and when, inferences derived from observed stimulus- and response-locked LPC effects differ.

This issue is also directly relevant to studies assessing ERP correlates of recognition memory across different experimental conditions or populations. For example, A diminished LPC effect has been used in clinical studies as an indicator for impaired memory retrieval in certain populations and when using pharmacological manipulations (Curran et al., 2006; Glaser et al., 2012; lakimova et al., 2005; Molnár et al., 2013; Rubin et al., 1999; Tóth et al., 2016). However, RTs also likely covary with experimental manipulations or participant characteristics of interest, such as age (Birren et al., 1980), diagnosis of Schizophrenia (Morrens et al., 2007) and pharmacological drug effects (e.g., midazolam, Thompson et al., 1999). This is not to suggest all previous research is invalided by the presence of the timing confound. Instead, we stress on the importance of clarifying whether changes in LPC effects could be partly attributed to the slower RTs, for example due to factors unrelated to the recollection process such as the non-decision time component specified in evidence accumulation models of decision-making (Brown & Heathcote, 2005; Ratcliff & Mcl<oon, 2008). We also note that RT differences are also relevant when investigating recognition memory using functional magnetic resonance imaging (Mumford et al., 2023; Yarkoni et al., 2009).

### 4.2. Other Contributions to LPC Effects

In addition to the response-locked LPC effect, we also identified other signals that could plausibly influence stimulus-locked measures of the LPC at left parietal electrodes. By contrasting hits and correct rejections, we found ERP differences shortly after the response (i.e., after a decision had been made), and a central midline, parietal stimulus-locked effect. We do not regard these as canonical LPC effects based on their timing and topography. While these novel findings warrant further replication, we suggest both are important to consider for investigation of LPC effects.

First, the post-decisional effect could similarly contribute to LPC effects in stimulus-locked ERPs, despite occurring after the recognition memory decision has been made. The timing of this post-decisional effect will also vary depending on the RT distributions in each condition of interest. This is important because LPC effects are typically assumed to reflect a recollection process that determines recognition memory decisions.

To our knowledge, no studies have investigated post-decisional effects on the LPC in recognition memory tasks (for a late frontal component, see Hayama et al., 2008; Johnson Jr et al., 1998). The interpretation of this post-decisional ERP effect is also limited in the current study because the vocal responses introduced significant and wide-spread electromyogenic (muscle) artefacts starting at the time of the response. It is also possible that this effect simply represents a carry-over of pre-decisional LPC effects. This again stresses the importance of using response-locked measurement of LPC to distinguish pre- and post-decisional effects.

Second, the stimulus-locked, central parietal effect could contribute to LPC effects in conventional stimulus-locked ERPs. This effect occurred over the time window typically used to measure the LPC, and temporally overlapped with the response-locked LPC effect. The overlap between the two neural effects could potentially account for previous descriptions of a bilateral LPC effect with a greater difference observed in left parietal areas (e.g., Kayser et al., 2007; Curran, 2004). Therefore, we emphasize the benefits of isolating the stimulus-locked subcomponents when investigating the response-locked LPC effect.

We identified a response-locked, midline parietal component that scaled in amplitude with decision confidence for both hits and correct rejections. This is consistent with previous observations of parietal, stimulus-locked ERPs varying with decision confidence in recognition memory tasks (Curran & Hancock, 2007; Woodruff et al., 2006). While the topography of the effect was distinct from the left-lateralised LPC effect, it nevertheless influenced amplitude measures at left parietal electrodes.

Our findings support the notion that LPC and confidence-related effects are (at least partially) distinct (Curran & Hancock, 2007; Woodruff et al., 2006). Notably, the topography of the confidence-related effects in our data is consistent with those recently reported in perceptual decision-making tasks using response-locked ERPs (Grogan et al., 2023; Ko et al., 2023), suggesting that we have identified a pre-decisional neural correlate of decision confidence that generalises across tasks. While recollection processes have been theorised to produce high confidence responses (Yonelinas, 1994), the effects of confidence at the midline parietal regions in our data do not appear to be simply a by-product of increased rates of recollection for trials with higher confidence ratings. Effects of confidence with the same topography were observed across both hits and correct rejections. In trials with correct rejections, recollection is not theorised to underlie high confidence responses.

Despite confidence-related effects showing a midline parietal locus, our analyses at electrode P3 demonstrate that effects of confidence can influence measures of the LPC at left parietal channels. This is particularly important when comparing LPC effects across conditions that vary with respect to reported confidence ratings. As higher confidence is also found to be associated with faster RTs (Ratcliff & Murdock, 1976; Robinson et al., 1997), effects of confidence may also intersect with effects on stimulus-locked ERPs relating to RT distributions as described above. With an understanding of these potential artefacts, in the stimulus-locked data, we observed similar patterns of amplitude changes with confidence shown in previous studies at left parietal regions (plotted in the Supplementary Material, Woroch & Gonsalves, 2010; Woodruff et al., 2006; Addante et al., 2012). Notably, variables that covary with memory performance are also theorised to be associated with decision confidence (Ratcliff & Stams, 2009; Reynolds et al., 2021). In order to better disentangle effects of confidence from hypothesised effects of recollection, it will be critical to delineate the LPC from confidence-related signals in future work.

### 4.4. FN400 Effects in Conventional ERPs and CSD-Transformed Data

We observed a typical FN400 effect when comparing hits and correct rejections in our standard ERPs. However, after CSD transformation this effect was no longer observable, and an effect was found with reversed polarity. This implies that the FN400 effect in our standard ERPs was likely due to volume conduction from more posterior areas. Using the same dataset, a reversed frontal effect could also be observed in Figure 2A from Weidemann and Kahana (2019), suggesting that our findings were not due to our data processing pipeline. Given the abundance of data, it is also unlikely that our results were due to noise or a lack of measurement precision. Our findings differ from prior studies using CSD transformations that reported FN400 effects at frontal electrodes (Johnson et al., 1998; Kayser et al., 2007; Kayser et al., 2009). As we did not observe an FN400 effect with a clear frontal source in our data, we did not perform further analyses using deconvolved ERPs.

### 4.5. Theoretical Implications

We propose that our finding of a response-locked LPC provides new perspectives for both single and dual process theories and interpretations. The later time window of LPC effects as compared to FN400 effects has been suggested to align with recollection processes that are slower than familiarity-related processes (Yonelinas, 1994, Yonelinas, 2002). However, we demonstrate this is not always the case for the left parietal LPC, as it can also occur with fast responses. Instead, we propose that the LPC effect represents a process that is time-varying, which precedes recognition memory decisions.

Under dual process theories, this is compatible with a recollection process whereby higher quality evidence is gathered for recognition memory decisions to terminate. To consider it as a time-varying process, the speed of recollection could be related to the internal noise of the recollection process that varies from moment to moment, and/or different degrees of total mnemonic information recollected from each attempt (i.e., across trials) with greater evidence producing faster and more accurate decisions. These two sources of variance that produce changes in RTs and performance have been assumed in Diffusion Decision Models (DDMs) to be captured by two separate parameters (Ratcliff & McKoon, 2008). However, despite intensive research into DDM and dual process theories, no study to our knowledge has tested dual process diffusion models of recognition memory. An investigation using such a model would provide an opportunity to test assumptions of the recollection process with regards to the observed RTs. For example, recollection can be specified as providing an either fixed amount of high-quality evidence assumed in classic dual process theories, or variable evidence conceptualised in dual process signal detection models (Onyper et al., 2010; Wixted, 2007).

Alternatively, the finding of a response-locked LPC is also compatible with the mnemonic accumulator theory of the LPC (Wagner et al., 2005) where memory-based evidence is accumulated and a decision is reached once the accumulation reaches a criterion. One important distinction regarding the accumulator theory is whether the accumulated evidence is limited to one process (e.g., mnemonic information) or an integration of processes supporting recognition memory (e.g., recollection and familiarity, Wixted, 2004). If LPC effects represent a single variable that supports recognition memory decisions, then the rate of change of the LPC waveform should scale with decision evidence. In models of retrieval, the strength of decision evidence can be calculated as the global similarity of the probe item to all the other target items. This global similarity could be computed externally based on item features (Osth et al., 2018) as well as internally based on the similarity in neural responses across items (e.g., Davis et al., 2014). Therefore, the relationship between computed global similarities and LPC signals could be tested empirically, and the results could provide insights into single process interpretations of LPC effects.

### 4.6. Limitations

Our findings should be interpreted with the following caveats in mind. First, our novel findings related to the LPC should be further replicated across other datasets and experiments. While the RIDE algorithm could successfully decompose ERP waveforms into subcomponents, this deconvolution technique relies on approximations and is not perfect. Future work could also compare results from the RIDE algorithm to other deconvolution methods such as algorithm employed in the Unfold toolbox (Ehinger & Dimigen, 2019).

Second, the functional role of the response-locked LPC needs to be further tested with other experimental manipulations and variables, with considerations of the RT and confidence-related issues discussed above. Although the dataset we analysed includes experimental manipulations that may help explain the function of the LPC signal, such as recall and encoding tasks, we are hesitant to draw specific inferences based on those manipulations. For example, the effect of previous recall on LPC effects may be difficult to delineate from greater confidence-related signals at central parietal electrodes in our data.

In addition, investigation on the post-decisional neural signals was also limited due to the artefacts introduced by speech when responding. Methods that better delineate these spatially overlapping neural effects, such principal component analysis (Haese & Czernochowski, 2022), could be used in future work.

### 4.7. Conclusion

By applying ERP deconvolution techniques to a large dataset recorded during a recognition memory task, we identified multiple sources of neural activity that contribute to classical LPC effects. Importantly, we show that LPC effects with a left-lateralised, parietal locus are time-locked to the response rather than stimulus onset. We also identified a response-locked neural correlate of decision confidence. Our findings highlight important issues that arise when interpreting existing LPC effects in stimulus-locked ERPs, and point to more accurate measurement practices using response-locked ERPs that can help avoid potential confounds. Our revised definition of the LPC and analytical approach could be adopted in future work to better characterise the functional significance of LPC effects in recognition memory paradigms.

## Data and Code Availability

The dataset is available upon request from the Computational Memory Lab at https://memory.psych.upenn.edu/Data_Request, which is a processed subset of the PEERS dataset published at https://0penneur0.0rg/datasets/ds004395/versi0ns/2.0.0. The MATLAB code for data pre-processing, and processed data to directly reproduce our results are available at https://osf.io/kur25.

## Author Contributions

Jie Sun: Conceptualization, Methodology, Software, Formal Analysis, Visualization, Writing — original draft, and Writing — review & editing. Adam F. Osth: Conceptualization, Project Administration, Supervision, and Writing — review & editing. Daniel Feuerriegel: Conceptualization, Project Administration, Supervision, Methodology, Visualization, and Writing — review and editing.

## Funding

This project was supported by an Australian Research Council Discovery Early Career Researcher Award to D.F. (ARC DE220101508) and the Graduate Research Scholarship awarded to J.S. from the University of Melbourne. Funding sources had no role in study design, data collection, analysis or interpretation of results.

## Declaration of Competing Interests

The Authors declare no competing interests.

## Acknowledgements

We would like to thank the Computational Memory Lab for generously sharing their data and Dr. Christoph T. Weidemann for prompt and helpful responses to questions related to the EEG dataset.

## Supplementary Material

Supplementary material for this article is available at https://osf.io/vafmt.

## Notes

https://osf.io/kur25/

### Competing Interest Statement

The authors have declared no competing interest.

