## Supplementary Material for "The Late Positive Event-Related Potential Component is Time-Locked to the Decision in Recognition Memory Tasks"

**Left Parietal ERPs for Trials with High and Low Confidence Ratings**

Studies that have investigated relationships between LPC amplitudes and decision confidence have often used four- or five-point confidence rating scales, and have defined high and low confidence categories for hits and correct rejections (Addante et al., 2012; Woodruff et al., 2006; Woroch & Gonsalves, 2010). To present ERPs in line with this analysis approach, we split our five-level confidence scale into either high (5) or low (1 to 4) confidence rating categories for hits. We included all confidence ratings for correct rejections.

We have plotted these ERPs at electrode P3 using stimulus-locked data in Figure 1A. ERPs were qualitatively more positive-going during a conventional LPC measurement window (400-800 ms) for high confidence hits compared to low confidence hits, and for high and low confidence hits compared to correct rejections. This is consistent with previous observations (Addante et al., 2012; Worch & Gonsalves, 2010).

We have also plotted ERPs using stimulus-locked ERPs whereby the stimulus-locked subcomponent had been subtracted from each trial prior to averaging (Figure 1B). In these ERPs, we observed qualitatively more positive-going ERP amplitudes for high confidence hits compared to correct rejections, however there were small to negligible differences between low confidence hits and correct rejections.

In addition, we have plotted response-locked ERPs with the stimulus-locked subcomponent subtracted from each trial in Figure 1C. Here, there are qualitatively more positive-going ERP amplitudes for high confidence hits compared to correct rejections. This difference appeared to extend into the time period after the initiation of the vocal response used to report recognition memory decisions (i.e., after the decision had been made).

Here, we note that the interpretation of these plots is complicated by issues relating to the response-locked nature of LPC effects, as well as co-occurring effects of confidence at electrode P3 and post-decisional ERP effects observed in the response-locked data. We present these plots to show that we have replicated typical patterns of ERP differences across high and low confidence hits and correct rejections as reported in prior work. We have further contextualised these patterns of ERP differences in our data by presenting both stimulus- and response-locked ERPs.

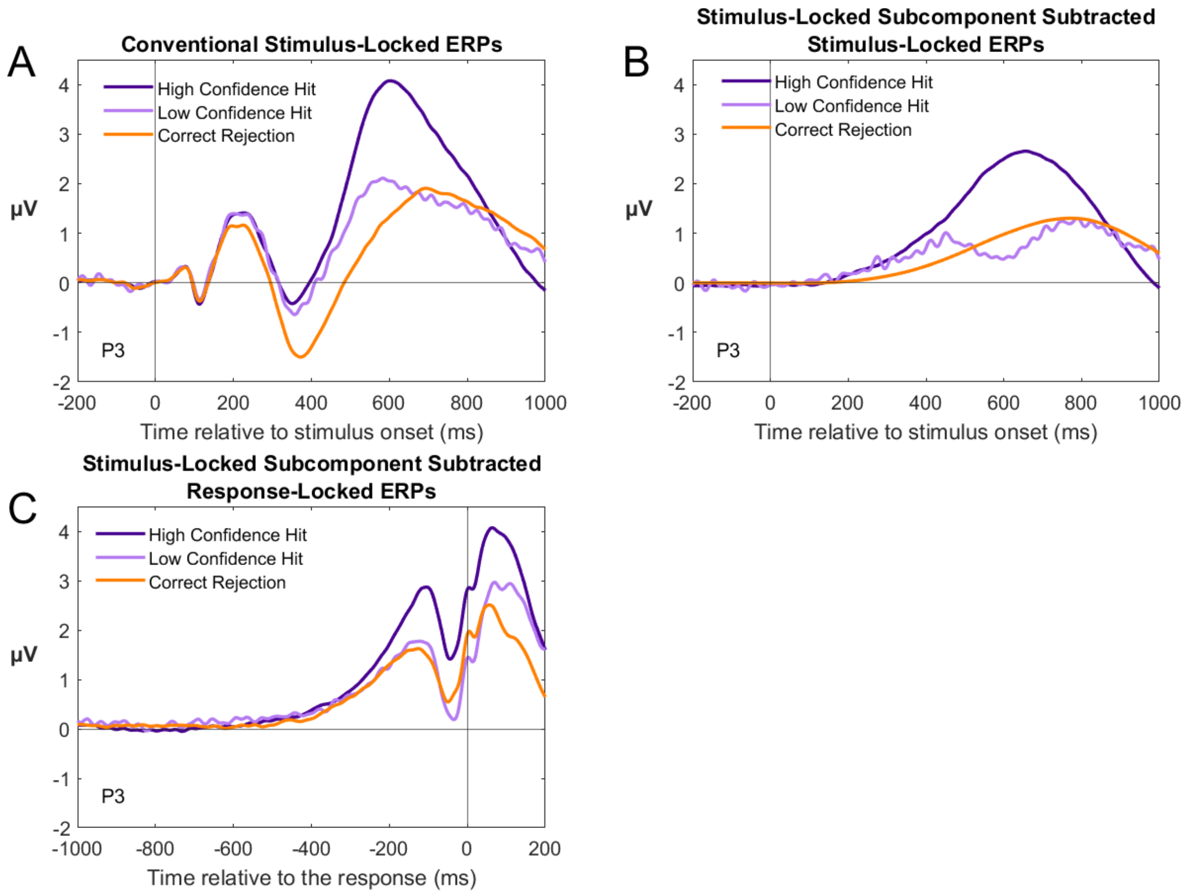


**Figure 1**. ERPs plotted by confidence category for hits and correct rejections at electrode P3. A) Conventional stimulus-locked ERPs. High confidence refers to a confidence rating of 5, and low confidence refers to confidence ratings of 1-4. B) Stimulus-locked ERPs with the stimulus-locked subcomponent subtracted. C) Response-locked ERPs with the stimulus-locked component subtracted.
